# Overexpression of the plastidal pseudo-protease *AtFtsHi3* confers drought tolerance without penalizing growth

**DOI:** 10.1101/2022.10.17.512522

**Authors:** Sam D. Cook, Laxmi S. Mishra, Hanna Isaksson, Isabella R. Straub, Miriam Abele, Sanatkumar Mishra, Christina Ludwig, Eric Libby, Christiane Funk

## Abstract

Drought is one of the most severe environmental stresses affecting plant biomass production and quality, however, the molecular mechanisms of drought response in plants remain unclear. Here, we demonstrate that overexpression of the Arabidopsis gene FTSHi3 under the influence of its endogenous, or the 35S constitutive promoter results in drought-tolerant phenotypes without penalising plant growth. FTSHi3 encodes a pseudo-protease located in the chloroplast envelope and knock-down mutants (*ftshi3-1*) have previously been found to be drought tolerant, but highly reduced in growth. Changes in FtsHi3 transcript abundance therefore seems to induce drought tolerance in *Arabidopsis thaliana*. Overexpression of FTSHi3 (*pFTSHi3-OE*) impacts leaf stomatal density, lowers stomatal conductance and increases water use efficiency. To explore the underlying mechanisms behind this, we compared the proteomes of *ftshi3-1* and *pFTSHi3-OE* to wild type plants grown under drought and watered conditions. Under drought conditions, the drought related processes ‘osmotic stress’, ‘water transport’ and ‘response to abscisic acid’ were enriched, indicating that *pFtsHi3-OE* and *ftshi3-1* mutants are more active in their response to drought than the wild-type. The proteins HSP90, HSP93 and TIC110 were more abundant in the knock-down mutant, which suggests that FtsHi3 might play a downstream role in chloroplast pre-protein import. Increased abundance of FtsH7/9 and FtsH11, FtsH12 and FtsHi4 in *ftshi3-1* combined with the fact that FtsH proteases function as homo- or heteromeric complexes suggests that these proteases may be possible interacting partners. To explore this, we constructed mathematical models that show FtsHi3 likely interacts with at least two other (pseudo-) proteases.

## Introduction

Drought is one of the most severe environmental stresses affecting plant biomass production and quality (Seleiman et al., 2021). Several classes of proteolytic enzymes have been shown to be involved in the response to drought (Vaseva et al., 2011; Fanourakis et al., 2020). For example, several cysteine proteases have been associated with drought in *Arabidopsis* and wheat (Botha et al., 2017). RNA sequencing studies on hexaploid sweet potato (*Ipomoea batatas*) showed higher expression levels of the ATP-dependent CLP proteases, whereas an aspartic protease in guard cells (*ASPARTIC PROTEASE IN GUARD CELL 1, ASPG1*) was significantly down-regulated in drought stress (Arisha et al., 2020). Overexpression of *ASPG1* also enhanced ABA (abscisic acid) sensitivity in Arabidopsis guard cells, reducing water loss and conferring drought resistance in *Arabidopsis* (Yao et al., 2012). Despite the knowledge of involvement of several proteases in adaptation to drought stress, the underlying molecular mechanisms are far from understood.

The membrane-bound FtsH (filamentation temperature-sensitive H) proteases are found in eubacteria, mitochondria and chloroplasts and are important for the degradation of cytosolic and membrane proteins (Smakowska et al., 2014; Nishimura et al., 2016; Bittner et al., 2017). As ATP-dependent metalloproteases, they contain a recognized AAA+ domain (IPR003593), a region of approximately 250 amino acids with the ATP-binding walker motifs (Walker A and B) and the second region of homology (SRH), which is positioned between the transmembrane helices and the proteolytic zinc-binding M41 peptidase domain (IPR000642) (Martin, 2021). Besides the proteolytically active FtsH proteases, pseudo-proteases, termed FtsHi (i for inactive (Sokolenko et al., 2002)), have been identified in the genomes of plants. Five FtsHi enzymes are present in *Arabidopsis thaliana*, all of which are localized in the chloroplast envelope. In these, the consensus sequence HEXXH of the Zn^+^-binding M41 peptidase domain is either absent (FtsHi3) or altered (FtsHi1/2/4/5) (Wagner et al., 2012). *Ycf*2 (hypothetical chloroplast open reading frame 2) has been suggested to encode a sixth FtsHi enzyme in *Arabidopsis thaliana* (Kikuchi et al., 2018), which also occurs in the green alga *Chlamydomonas reinhardtii* (Ramundo et al., 2020). FtsHi1, 2, 4, 5 form an import motor complex with FtsH12, a NAD-dependent malate dehydrogenase (MDH) (Schreier et al., 2018, Mielke et al., 2020) and possibly Ycf2 (Kikuchi et al., 2018). Homozygous mutants lacking functional FtsHi1, 2, 4, or 5 also show a seed-lethal phenotype (Kadirjan-Kalbach et al., 2012; Lu et al., 2014; Wang et al., 2018; Mishra et al., 2019), whereas heterozygous *FTSHi* mutants and plants carrying point/missense or “*weak*” mutations display a pale-green-seedling phenotype. These plants have smaller rosettes throughout their life span (Kadirjan-Kalbach et al., 2012; Lu et al., 2014; Mishra et al., 2019; Mishra et al., 2021) and often contain variegated leaves (Wang et al., 2018). FtsHi3 was not identified as a component in the FtsH12/FtsHi1,2,4,5/MDH/Ycf2 complex (Kikuchi et al., 2018), and its interacting partners remain unknown. Also, homozygous *ftshi3* mutants are not embryo lethal (Kikuchi et al., 2018; Mishra et al., 2019; Mishra et al., 2021). Recently, the *ftshi3-1* knockdown mutant, which has significantly reduced *FTSHi*3 expression (termed *ftshi3-1* (kd) in Mishra et al., 2021), was shown to be drought tolerant. However, its growth was strongly affected, and seed germination was somewhat delayed.

Here, we show that overexpression of the *FTSHi3* gene, results in a drought-tolerant phenotype without the reductions in growth seen in *ftshi3-1*. We demonstrate that overexpression of *FTSHi*3 impacts leaf stomatal density, lowers stomatal conductance and increases the water use efficiency index (WUEi). The levels of several ABA-responsive genes in *pFtsHi3-OE* plants were elevated (relative to the wild type (WT)) in watered conditions, although, their expression in drought did not change substantially in these lines. Comparing the proteomes of *ftshi3-1* and *pFtsHi3-OE* to WT revealed that several drought associated processes are repressed in the mutants, such as response to water deprivation and response to abscisic acid (ABA). The chloroplast envelope located AtFtsH7/9 and AtFtsH11, AtFtsH12 and AtFtsHi4 were detected in *ftshi3-1,* independent of the growth conditions, but not in *pFtsHi3-OE* or WT and therefore might be possible complex partners substituting for the lost subunit. Mathematical modeling was performed to understand how variation in the transcript abundance of *FTSHi3* can on one hand lead to drought tolerance in both overexpression and mutant lines yet affects plant growth so differently.

## Results

### Characterization of *Arabidopsis* plants with elevated expression of *AtFtshi3*

Previously, we characterized a knock-down mutant in Arabidopsis (*ftshi3-1*) with significantly reduced *FTSHi*3 expression (0.1% of WT). This mutant is considerably smaller than WT and displays a chlorotic phenotype throughout its lifetime (Mishra et al., 2021). Plants depleted of *FtsHi3* are also, unexpectedly, drought-tolerant compared to WT (Mishra et al., 2021). To understand the role of FtsHi3 in overall plant development, we generated lines expressing *FTSHi3* under the control of its endogenous promoter (*pFtsHi3-OE*) (Figure 1), or of the constitutive *35S* promotor (*35SFtsHi3-OE*) (Figure S1)*. FTSHi3* transcript levels were determined in *pFtsHi3-OE* lines 1-10 (Figure 1C) and *35SFtsHi3-OE* lines 5-8 (Figure S1C). The *pFtsHi3-OE* lines 1-5 showed approximately 6-to 8-fold expression of *FTSHi3,* while lines 6-8 showed 4-fold increases relative to the WT. *FTSHI3* expression was elevated two-fold in *pFtsHi3-*OE lines 9 and 10. Transcripts in the *35SFtsHi-OE* lines were typically twice that of WT (Figure S1C). Based on these results, we selected two representative lines (*pFtsHi3-OE1* and *pFtsHi3-OE2*) for additional physiological studies (Figure 1D).

**Figure 1:**
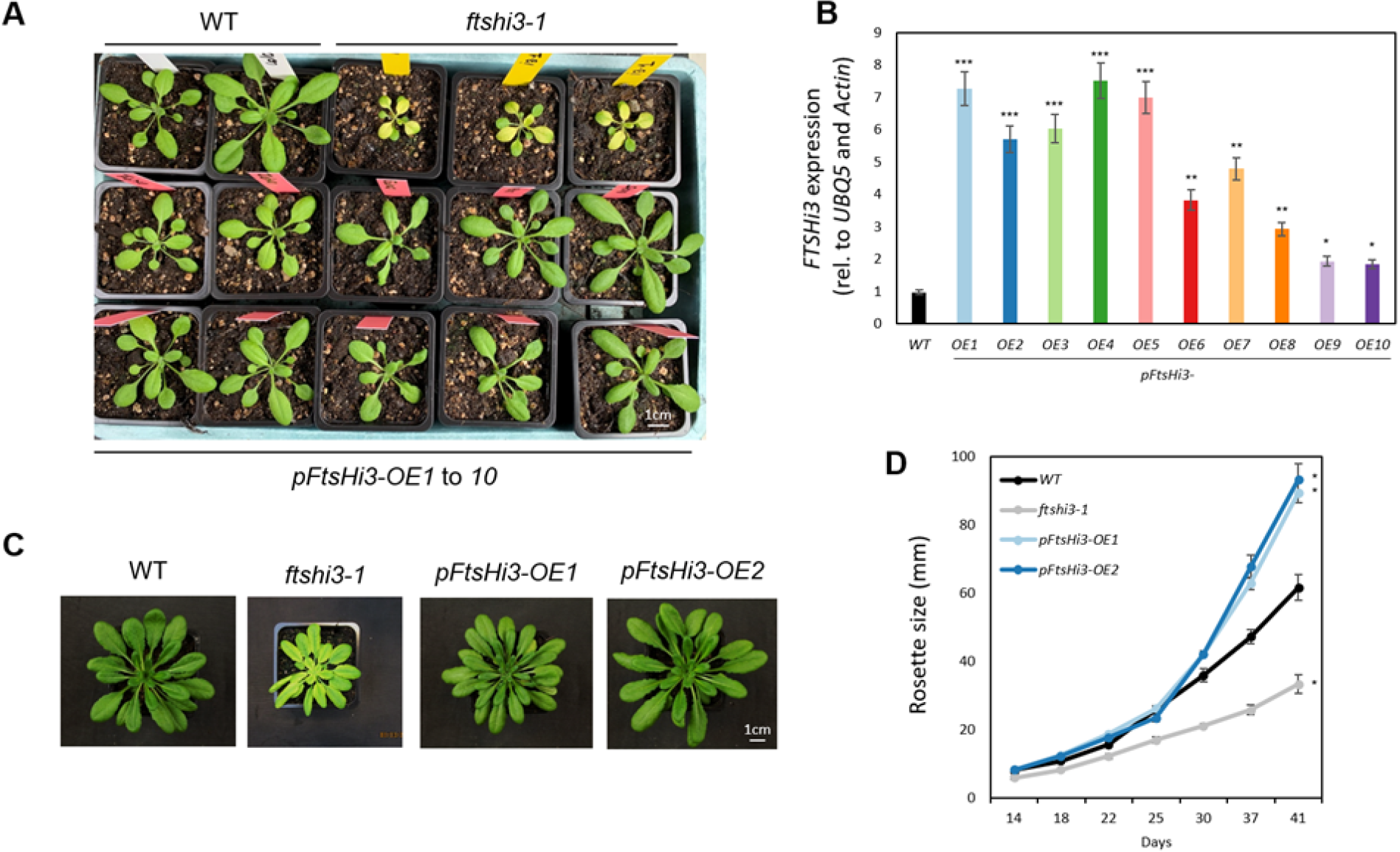
**A,** Phenotypes of WT (top left, white labels), *ftshi3-1* (top right, yellow labels) and *pFtsHi3-OE1* to *-OE5* (middle, red labels) and pFtsHi3-OE6 to -OE10 (lower, red labels) at 3 weeks of age under long day (LD) conditions (16/8 hr D/N cycle, 22°C). **B,** Expression of *FTSHi3* relative to *UBQ5* and *actin* in each of the *pFtsHi3-OE* lines at 3 weeks of growth under short day (SD), soil grown conditions (n=3). **C,** Phenotypes of WT, *ftshi3-1* and the two representative lines (*pFtsHi3-OE1* and *-OE2*) at 6 weeks of age under SD conditions (8/16 hr D/N cycle, 22°C). **D,** Rosette diameter of WT, *ftshi3-1*, *pFtsHi3-OE1* and *-OE2* under the same conditions as **C** (n=8). Data are means ± standard error, asterisks indicate significance; p<0.05 (*), p< 0.01 (**), p< 0.001 (***) and scale bars are as indicated.

Under short day conditions, which inhibit bolting, both representative lines appeared similar to the WT as seedlings, but their rosettes were significantly larger at later stages (Figure 1D, p<0.05). The rosettes of *35SFtsHi3-OE* lines also grew to a larger size at later time points (Figure S1B). A similar growth pattern was observed in all lines for both constructs under long day conditions (data not shown). We further observed significantly larger root growth (around 20%) (Figure S2A-B) and a trend towards increased number of lateral roots in *pFtsHi3-OE1* and *pFtsHi3-OE2* (Figure S2C). Contrary to *ftshi3-1* (Mishra et al., 2021), chloroplasts in the first true leaves of *pFtsHi3-OE1* and *pFtsHi3-OE2* lines display fully developed ultrastructures (Figure S2D), have increased width and an elevated number of thylakoid membranes (with a significantly higher degree of grana stacking) (Figure S2E-F).

### Overexpression of *FTSHi3* confers drought tolerance without penalizing plant growth

The *ftshi3-1* mutant has significantly reduced *FTSHi3* transcripts, which result in drought tolerance (Mishra et al., 2021). Here we investigated the response to drought stress in the 10 *pFtsHi3-OE* and the 4 *35SFtsHi-OE* lines (Figure 2, 3, Figure S3). Prior to droughting, the rosette size and phenotypes of the OE lines were generally consistent with the WT. This was retained throughout the 15-day drought period, where despite cessations in growth, most *pFtsHi3-OE* lines had larger rosettes and higher final biomass (Figure 2A, C). Curiously, all *35SFtsHi3-OE* lines continued to grow throughout the drought period (Figure S3A), resulting in even higher final dry weights (Figure S3C).

**Figure 2:**
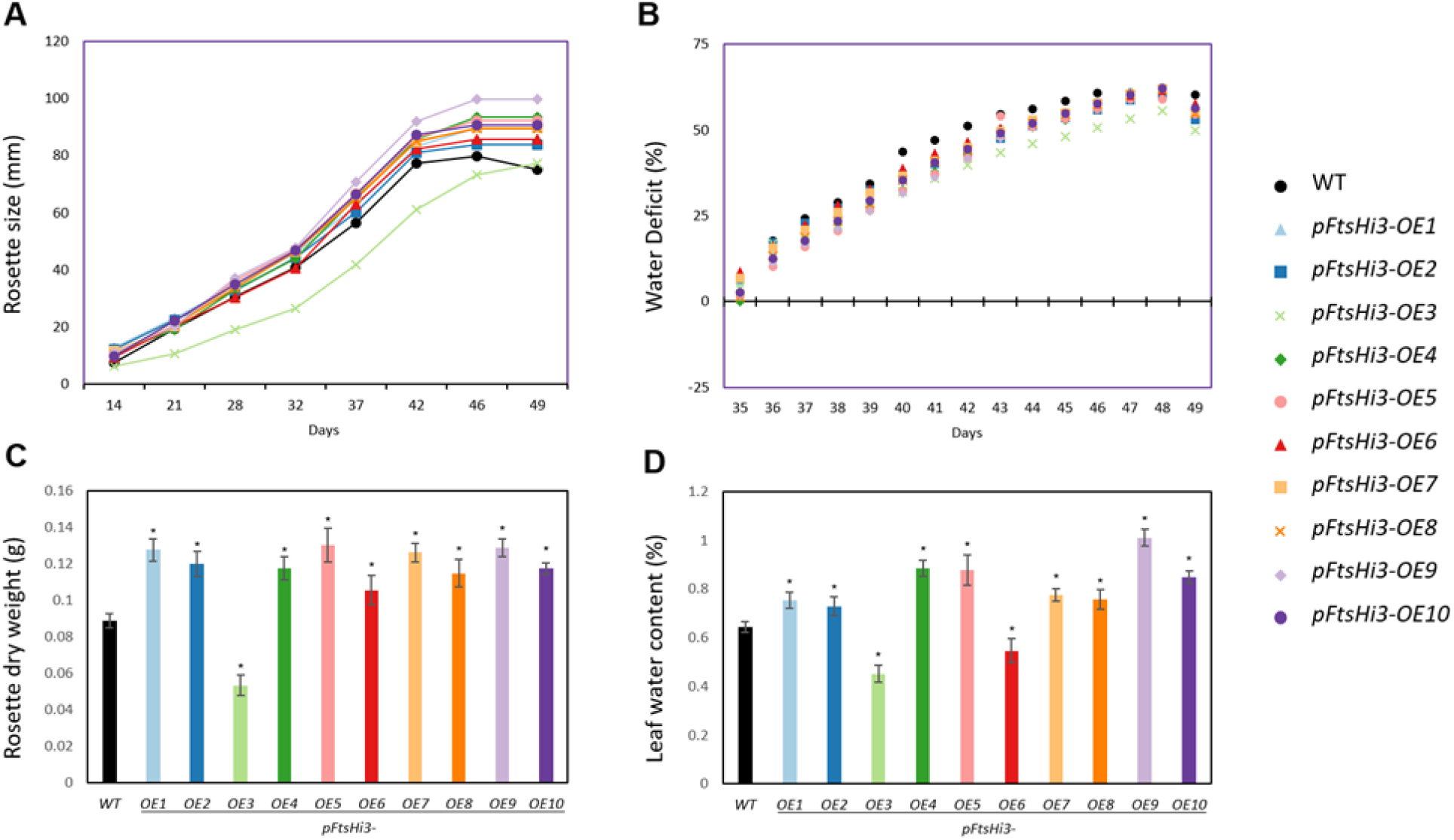
Effect of drought on WT and *pFtsHi3-OE1* to *OE10* lines grown in a growth chamber under SD, soil grown conditions (8/16 hr D/N cycle, 22°C). **A,** Rosette diameter prior to, and following, drought induction at day 37. **B,** Progressive % water deficit during drought. **C,** Dry weight of harvested rosettes following drought. **D,** Calculated leaf water content %: (Biomass of watered control) - (Biomass of droughted) / (Biomass of watered control). Data are means ± standard error (n=15), significances as per Figure 1.

**Figure 3.**
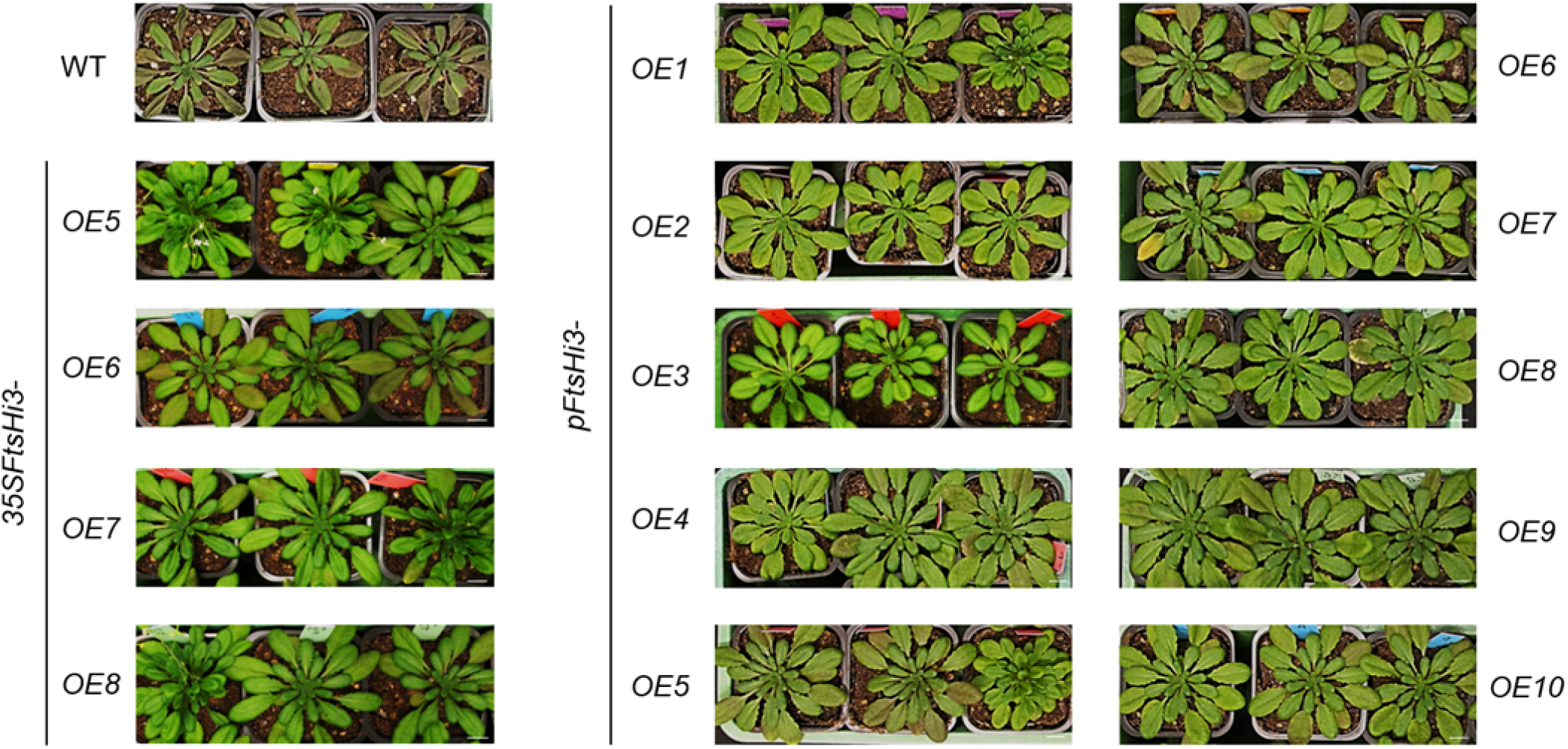
Representative post-drought phenotypes of WT, *35SFtsHi3-OE5-8*, and *pFtsHi3-OE1-10* transgenic plants exposed to drought stress for 15 days under SD conditions (8/16 hr D/N cycle, 22°C). Plants were germinated on plates containing MS + 1% sucrose and transferred to pots after 7 days. Following ∼4 weeks of growth (30 days), plants were droughted and images were taken at ∼7 weeks of age.

Interestingly, we noted that the degree of wilting was significantly reduced in all OE lines (Table S1, Figure 3). This strongly correlated with the survivability of the different lines following drought stress. The extent of water deficit during progressive drought was similar between WT and the *pFtsHi3-OE* lines (Figure 2B), although the WT appeared to have a higher deficit. The *35SFtsHi3-OE* lines were much more variable and were more deficient at the end of the experiment (Figure S3B). The leaf water content of most overexpression lines (except for *pFtsHi3-OE3* and *-OE6*) was higher than that of WT plants (Figure 2D, Figure S3D).

### Gas exchange qualities, stomatal physiology and ABA response are altered in *pFtsHi3-OE* lines

Further characterization of the impact of elevated *AtFTSHi3* expression on drought stress was carried out using the two representative lines *pFtsHi3-OE1* and *-OE2* as well as *35SFtsHi3-OE5, -OE7 and -OE8*. Gas exchange parameters were determined immediately before and following 14 days of drought (Table S2). Prior to drought, net photosynthesis (*A_N_*) and stomatal conductance (*gs*) were variable across all genotypes, while CO_2_ concentration inside the leaf (*Ci*) was consistent. After 14 days of drought treatment, the WT displayed significantly lower *A_N_* and *gs* compared to the OE lines (Table S2). Interestingly, the *Ci* values following drought were elevated in the WT but decreased in the OE lines. The water use efficiency (WUE intrinsic), i.e. the ratio *A_N_/ gs*, was significantly increased in all overexpression lines following drought treatment (Table S2).

The levels of abscisic acid (ABA) were investigated pre- and post-drought in the WT and *pFtsHi3-OE* representative lines. While the ABA levels in pre-drought conditions were similar in all genotypes (Figure S4C), the response of the pFtsHi3-OE lines was subdued following drought (∼2-fold increase) when compared to the WT (∼6-fold increase, Figure S4C). To understand these findings, we compared stomatal density and size in WT and *pFtsHi3-OE,* and *pFtsHi3-OE2* lines. WT plants contained an average of 252 stomata per mm^2^ compared to 200 and 204 in *pFtsHi3-OE1* and *pFtsHi3-OE2* lines, respectively (Figure S4B). In addition to the reduced number of stomata, their average size (width to length) was also reduced in the OE lines (18.9 by 14.9 μm for WT, 17.5 by 13.9 μm for *pFtsHi3-OE1* and 16.7 by 12.1 μm for *pFtsHi3-OE2*). We further tested stomatal closure with treatment of 10 μM exogenous ABA, although all lines responded similarly (Figure S4 D, E).

As ABA was reduced in *pFtsHi3-OE1* and *pFtsHi3-OE2* under drought conditions, we suspected that the expression of ABA-responsive genes might also be altered. Thus, we evaluated the expression patterns of nine ABA-responsive genes under drought and watered conditions (Table S3). The marker genes: *Responsive-to-desiccation 29A, 29B*, and *22* (*RD29A, RD29B, RD22*), *Dehydration-responsive-element-binding-protein 1A* and *2A* (*DREB1A, DREB2A*), *9-cis-epoxycarotenoid-dioxygenase* (*NCED3*), *Cold-regulated 47* (*COR47*) and *Drought-induced 21* (*DI21*) all displayed consistently higher expression in the OE lines under watered conditions. While in drought, *RD29B, DREB2A and COR47* were all highly elevated in the WT, but non-responsive in the OE lines. This is consistent with the muted accumulation of endogenous ABA under drought and suggests that the OE plants are already in a perceived state of drought under watered conditions, even though they contain WT levels of ABA. A similar pattern was observed in *Drought-induced 21* (*DI21*), which was slightly upregulated in the OE lines, but highly expressed in the WT under drought conditions. Curiously, RD22 displayed reduced expression in the WT upon entering drought, even though its expression in the OE lines was significantly increased.

### Plants with modulated FtsHi3 content show changes in proteome composition during water deficit

Changes in abundance of FtsHi3 seem to induce drought tolerance in *Arabidopsis thaliana*. However, while the absence of FtsHi3 (*ftshi3-1*) additionally leads to a chlorotic, dwarf phenotype (Mishra et al., 2021), overexpression of this pseudo-protease resulted in plants that are phenotypically similar to, or larger than the WT. To explore the effect of FtsHi3 on drought tolerance, we compared the proteomes of *ftshi3-1*, *pFtsHi3-OE1* and WT leaves under both watered and drought conditions. Using an S0 threshold of 2 (Effectively minimum fold change) and a false discovery rate (FDR) adjusted t test p-value cutoff of 0.01, we observed a number of proteins with altered abundance in selected comparisons; *ftshi3-1* vs WT under drought conditions: 762, *ftshi3-1* vs WT under watered conditions: 140, and *pFtsHi3-OE1* vs WT under drought conditions: 75 (Figure S5A-C). Curiously, there were no differentially abundant peptides in the watered *pFtsHi3-OE1* vs WT comparison (Figure S5D), indicating no clear functional difference in protein composition of WT and *pFtsHi3-OE1* lines under watered conditions. Surprisingly, although being overexpressed in *pFtsHi3-OE1*, we were unable to detect any peptide corresponding to FtsHi3.

The differentially abundant protein lists were further compared to segregate AGI’s (Arabidopsis Genome Initiative) and identify elements common to major contrasts (Figure S5). Most (64 of 75) differentially abundant proteins from the *pFtsHi3-OE1* vs WT comparison in drought condition (Figure S5C) were also detected in the *ftshi3-1 vs WT comparison* (Fig S6A). 97 AGIs were shared by the *ftshi3-1*: WT comparisons in watered and drought conditions. However, only seven AGIs were shared between the droughted *pFtsHi3-OE1*: WT and the watered *ftshi3-1*: WT contrasts. These seven AGIs include ribosomal proteins (ATCG01120, AT3G53740, AT2G37600, AT4G15000), a protein responsible for cell wall integrity (AT5G66090), a mitochondrial protein of the adenylate kinase family (AT5G50370) and an NADH dehydrogenase family protein (ATCG01100).

Given that the *pFtsHi3-OE* lines appeared phenotypically similar to the WT in growth yet still displayed a drought-tolerant phenotype, we compared the lists of altered AGIs to explore biological processes that might be controlling these factors. We separated out AGIs that were specifically altered in *ftshi3-1* (relative to the WT) to identify growth-related processes, and common AGIs (to *ftshi3-1* and *pFtsHi3-OE1* relative to the WT) to explore drought resistance (Figure S5E). We subsequently conducted ontological analyses on these AGI lists to highlight the enrichment of biological processes in our data sets (Tables S4-6). In the AGI list of peptides elevated only in the *ftshi3-1* mutant (under drought conditions), several processes associated with photosynthesis in general are enriched (Table S4). Protoporphyrinogen IX biosynthesis, chlorophyll biosynthesis, chloroplast rRNA processing, chloroplast protein import, thylakoid membrane organization and chloroplast organization are all over-represented. Additionally, many isoprenoid metabolic processes (geranylgeranyl diphosphate biosynthesis, geranyl diphosphate metabolism and farnesyl diphosphate biosynthesis) and amino acid biosynthetic processes (Valine, Isoleucine, Leucine, Lysine biosynthesis) were also enriched in *ftshi3-1*. We found that inositol biosynthesis, anthocyanin-containing compound biosynthesis and Phenylalanine catabolism were all highly enriched (>100-, 96.56- and 86.91-fold, respectively) in the repressed, *ftshi3-1* specific, AGI list (Table S5). Curiously, we also observed repression of several drought associated processes, such as response to desiccation and response to abscisic acid (ABA), implying that *ftshi3* mutant plants appear, at least partially, to be repressing drought response, relative to the WT.

To visualise the different ontological processes identified above, we constructed STRING plots (string-db.org) to explore known protein-protein interactions within the AGI lists specific for the *ftshi3-1* line. We observed three large clusters in the elevated protein list (Figure S6, High confidence interactions, MCL, Inf = 2) corresponding to tetrapyrrole (chlorophyll and heme) biosynthesis, chloroplast organisation and RNA processing. We also saw several smaller clusters that included protein trafficking enzymes (TICs and TOCs) as well as affiliated chaparone elements (CR88 and HSP70). In the repressed AGI STRING plot (Figure 4 we observed clusters of known drought/ cold responsive proteins (such as LEA14 and COR47), flavonol biosynthesis proteins (e.g. TT4/TT5) and photosyntetic proteins (e.g. the PsbQ-like proteins PnsL2/3) as well as a distinct cluster containing all three inositol 3-phosphate synthase proteins, which catalyse the rate limiting step of myo-inositol biosynthesis (Donahue et al. 2010).

**Figure 4.**
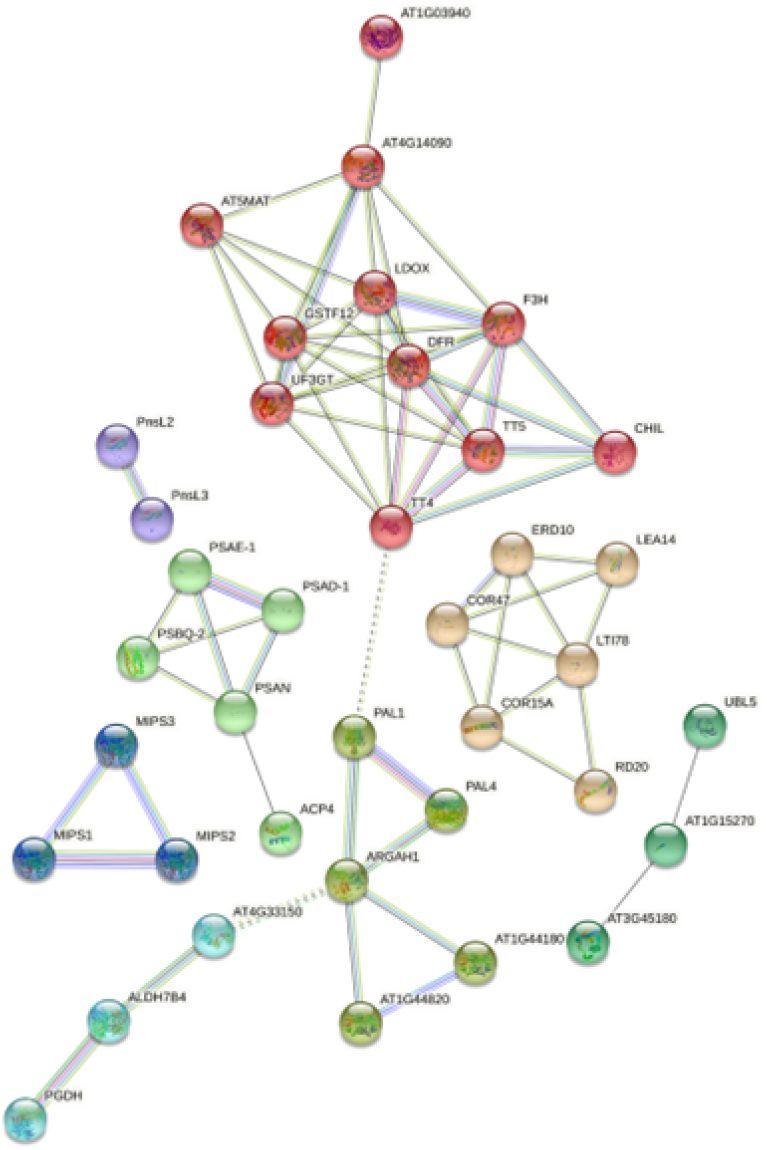
STRING plot showing protein: protein interactions between proteins **repressed** in *ftshi3-1* relative to WT under drought conditions. Data were filtered to retain only high confidence interactions (0.700) and disconnected nodes were removed. The resulting elements were clustered using an MCL inflation parameter of 2. Clusters are distinguished by node color and line color indicates number and type of evidence for interactions (see string-db.org/), dashed lines indicate borders of clusters. Protein names or AGIs are provided adjacent to nodes.

In the ontological analysis of AGIs common to *ftshi3-1* and *pFtsHi3-OE1* (Table S6), enzymes involved in proton transmembrane transport and response to water deprivation were highly enriched (43-and 21-fold, respectively). Additionally, the processes; response to cold, response to abscisic acid and response to osmotic stress were also strongly enriched. Given that the osmotic stress related processes are composed of AGIs that are repressed in both the *pFtsHi3-OE1* and *ftshi3-1* contrasts, this suggests that these processes may be responsible for conferring drought resistance to the *pFtsHi3-OE1* and *ftshi3-1* lines. Further exploration of these groups reveals quite a small number of AGIs that are present in several of these categories. To visualize their relative abundances (based on log LFQ values), we constructed a heatmap to compare the droughted WT, *pFtsHi3-OE1* and *ftshi3-1* lines as well as the watered WT (Figure 5). The abundances of the ABA inducible KIN1, the AKR4C9 aldo-keto reductase, the ABCG30/36/37 transporters and the Late Embryogenesis Abundant protein LEA4-5 appeared juxtaposed between the WT and the *pFtsHi3-OE1/ ftshi3-1* lines, while the abundance of the Thiamine biosynthesis protein THI1 abundance was increased more so in *ftshi3-1* than *pFtsHi3-OE1*. Relative to the droughted WT, we also found protein abundance of the putative chitinase CHI, LEA7, a lipid transfer protein: LTP4, AKR4C8, a drought responsive protein LTI65 (RD29B), a GTPase RAB18, and alcohol dehydrogenase ADH1 to be repressed in both *ftshi3-1* and *pFtsHi3-OE1*, but this was obviously more severe in the OE line.

**Figure 5.**
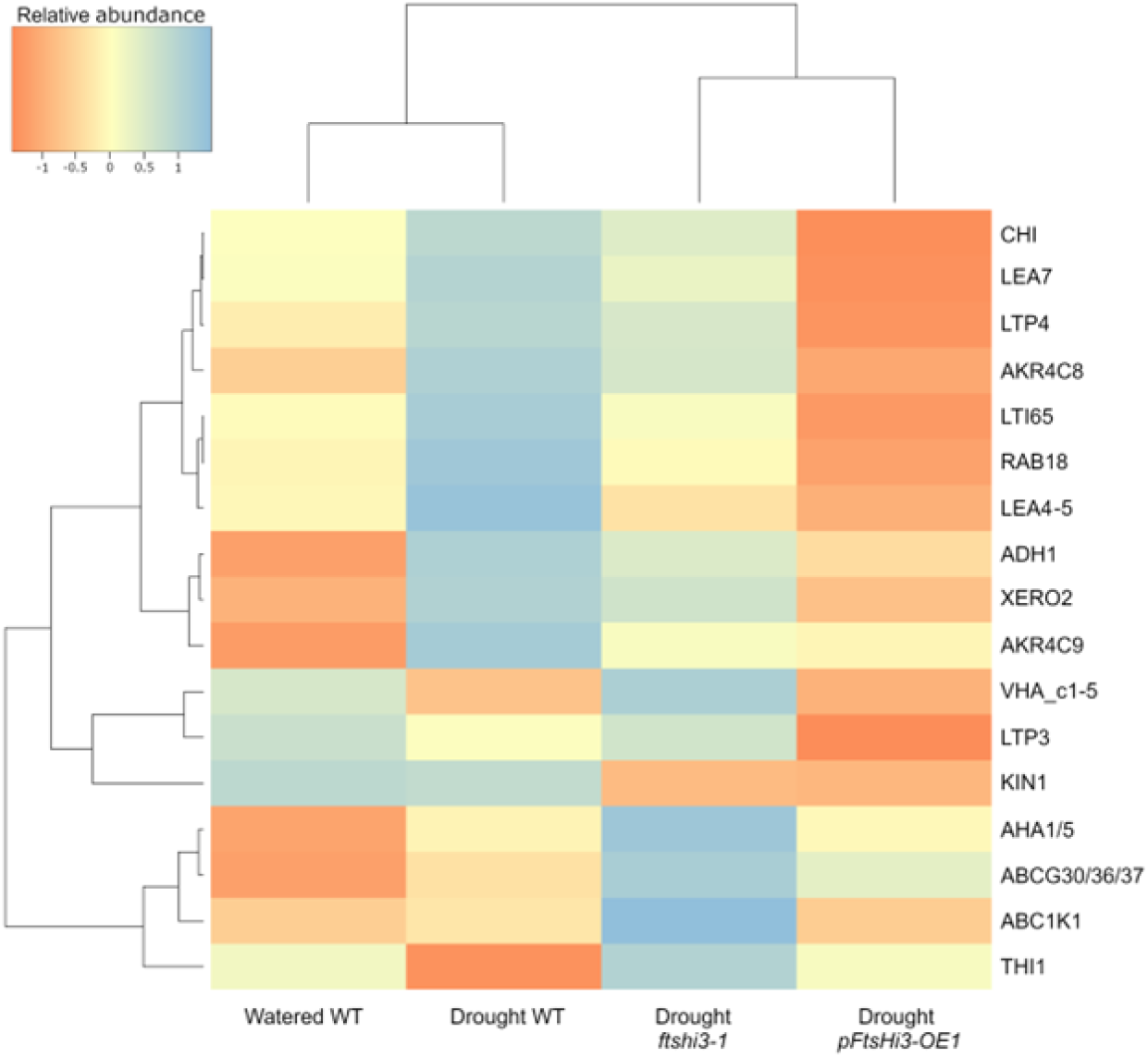
Heatmap showing mass spectrometric intensities (log_10_ LFQ) of AGIs that were differentially abundant in both *pFtsHi3-OE1* and *ftshi3-1* lines relative to the WT under drought conditions (S0 >2, p-value <0.01). Dendrograms represent dissimilarities between contrasts (above) and proteins (left).

### Hypothetical interacting partners

FtsH proteases are known to form homo- or heteromeric complexes (Sakamoto et al., 2003). Curiously, in the proteomic data we observed several interesting trends amongst the greater FtsH family with regards to peptide abundance in different genotype: treatment comparisons (Figure S7). Abundance of the FtsH proteases located in the chloroplast thylakoid did not vary between the different lines and treatments, which suggests that these elements are not particularly responsive to drought, and do not appear to be influenced by presence or absence of a functional FtsHi3. The chloroplast envelope located AtFtsH7 and AtFtsH9 (which are indistinguishable by mass spectrometry because only peptides shared between both proteins were detected), AtFtsH11, AtFtsH12 and AtFtsHi4, however, were present only in the *ftshi3-1* (under both conditions). In contrast to this, the mitochondrial FtsH4 was only detectable in drought conditions and only in the WT and *ftshi3-1* lines. Finally, FtsH3 and/ or FtsH10 (which also share a common identifying peptide) were detectable in all comparisons, bar the watered WT.

A variety of homo- or heteromeric complexes would provide a possible mechanism that could explain how differential expression of the FtsHi3 pseudo-protease induces characteristically different phenotypes: decreasing growth rate with lower expression (monotonic behavior) and increasing drought resistance with lower as well as higher expression (non-monotonic behavior). We therefore created a mathematical model to identify the minimal number and types of interactions between FtsHi3 and possible complex partners and/or substitutes (AtFtsH7 and/or AtFtsH9, AtFtsH11, AtFtsH12 and AtFtsHi4 (based on Figure S7)) that would recapitulate our experimental observations. The model was based on the following simplifying assumptions to produce a more general model: Each heteromeric structure is formed by only two types of homo-trimers in specific stoichiometric ratios of 1:1. We also assumed that both growth rate and drought resistance are determined by the relative abundance of different heteromeric complexes - thus it is not just the absolute amount of a specific protease, but its proportion in the membrane. We then considered a range of different biochemical models with varying numbers of FtsHi/FtsH homo-trimer subunits forming hexamers (3 or 4, less than 3 could not generate the behavior) and types of interactions (see Methods). Using a random sampling method we identified, which models (number of protease subunits and types of interactions) are most likely to reproduce our experimental findings (Figure 6).

**Figure 6.**
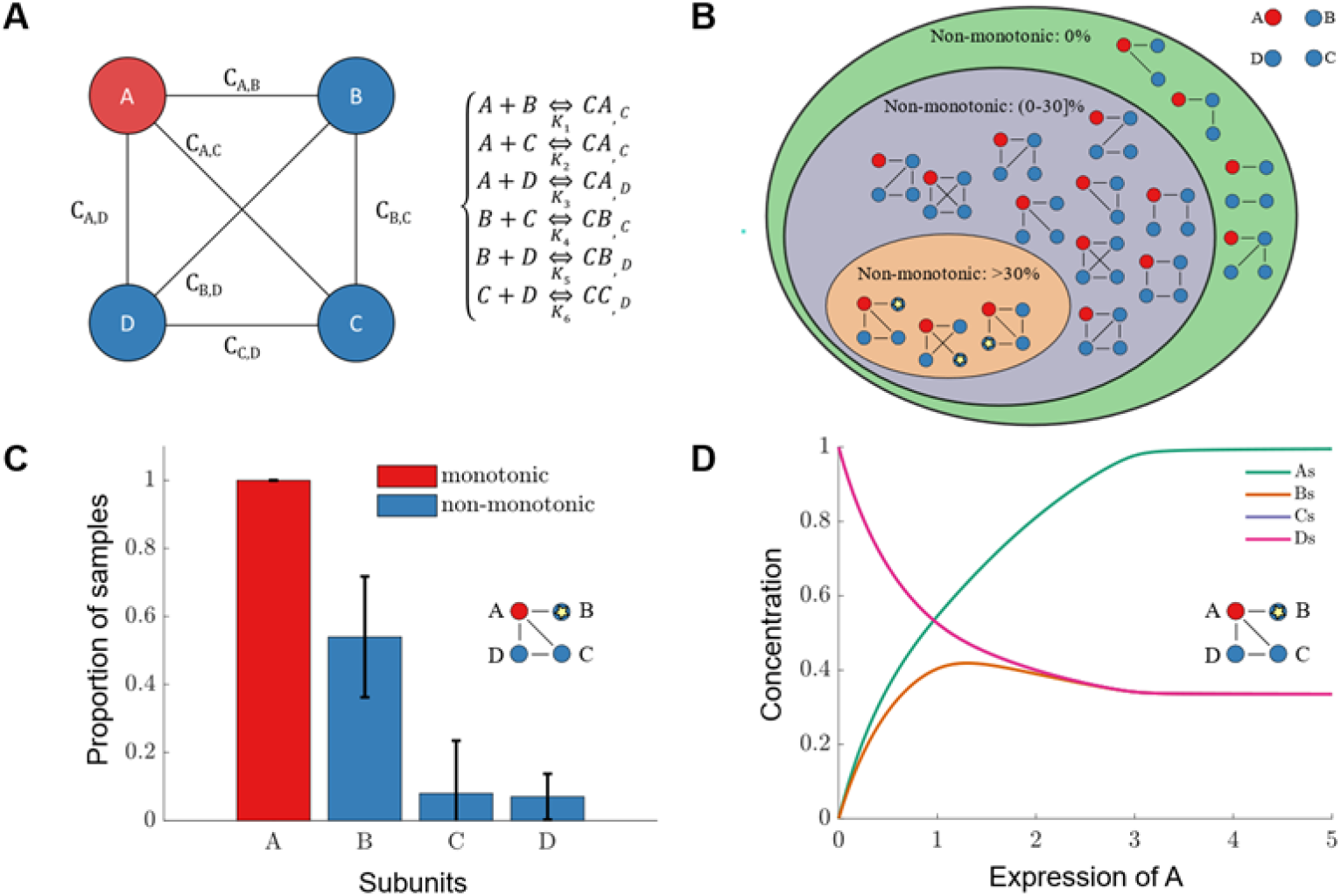
Mathematical modelling to explain the phenotypic variation concerning growth and drought tolerance of *FTSHi3* mutant lines. **A,** Schematic showing the possible interactions between (Pseudo-) protease homotrimers in the formation of the putative *FtsHi3* hexamer (based on our simplifying assumptions). **B,** Interaction diagrams of all possible binding partners in systems with 3 or 4 different subunits, organized by the frequency in which we find non-monotonic behavior. Red circles indicate subunit A, the FtsHi3 pseudo-protease homotrimer, and blue circles show other possible subunits B, C and D (based on our proteomic data: FtsH7 and/or 9, FtsH11, FtsH12, FtsHi4). Stars indicate the subunit with the most non-monotonic behavior. **C,** Bar graph showing the proportion of samples (mean of 10 samples of 100 simulations) that have monotonic behavior for subunit A (red) and non-monotonic behavior for other subunits (blue). The most isolated subunit (with star) is the one with most non-monotonic behavior. **D** Concentrations of complexes containing different subunits (based on the interaction diagram C) are shown as a function of increasing expression of subunit A (*FtsHi3*). The arch in complexes containing subunit B drive the non-monotonic behavior.

All models could recapitulate the monotonic behavior of growth because complexes that involved the FtsHi3 pseudo-protease naturally increased with increasing expression of FtsHi3. In contrast, the non-monotonic behavior of drought tolerance was less often observed in our models (Figure 6B). A common feature of those models with non-monotonic behavior was that FtsH proteases, especially FtsHi3, interact with multiple partners. Allowing different proteases to belong to several different heteromeric complexes creates opportunities for competitive binding that could shift the proportions of complexes (see Figure 6C, D). In models where FtsHi3 had a single specific partner, we did not observe non-monotonic behavior. We explored the robustness of the findings of our mathematical model for different parameter regimes by studying the dynamics for all 17 possible interactions for complexes including 3 or 4 subunits (Figure S8). The reader should note that the interactions in panels B) and L) stand out with having zero detected monotonic behavior. In these interactions all complexes contain the ‘**A**’ homo-trimer, therefore the proportion of the ‘**A**’ subunit equals 1 for each line (WT, *ftshi3-1* or *pFtsHi3-OE1*). Apart from these two special cases, monotonic behavior is detected in all other interactions. Non-monotonic behavior is more commonly found in interactions where compound A could bind to more than one other subunit (in Figure S8 panels D-G should be compared to panels L-Q). : In general, we observed that the models found in this paper were those, in which FtsHi3 could interact with at least two other proteases.

## Discussion

### FtsHi3 overexpression results in a drought tolerant physiology

Despite their proteolytic inactivity, FtsHi pseudo-proteases have been shown to be directly or indirectly involved in plant stress responses (Kadirjan-Kalbach et al., 2012; Lu et al., 2014; Kikuchi et al., 2018; Wang et al., 2018; Mishra et al., 2019; Mishra et al., 2021). While FtsHi1,2,4 and 5 are known to be subunits of a complex functioning in protein import into the chloroplast (Kikuchi et al., 2018; Schreier et al., 2018), the function of FtsHi3 remains enigmatic. The *ftshi3* knock-down mutants (Kikuchi et al., 2018; Mishra et al., 2019; Mishra et al., 2021) as well as the overexpression lines described here result in plants that are drought tolerant. However, contrary to *ftshi3* knock-down mutants, the *FTSHi3* overexpression lines displayed enhanced growth at later stages of plant development (Figure 1, Figure S1). This is mirrored in our drought experiments, where the dry rosette biomass was significantly higher in *pFtsHi3-OE* lines (Figure 2C). Curiously, despite having intermediate *FtsHi3* expression, the 35SFtsHi3-OE lines continued to grow following drought induction, while the *pFtsHi3-OE* lines did not (Figure S1C, Figure S3). This is remarkable as stress tolerant overexpression lines often display growth retardation when driven by a 35S promoter, yet are less severely affected by native promoter regions (Kasuga et al., 1999; Yi et al., 2010). The same trend is observed here (i.e. weakening of effect under native promotion) and can generally be attributed to the recognition and presence of the regulatory elements present in the endogenous promoter sequences (Hernandez-Garcia and Finer, 2014).

Although the overexpression lines displayed similar or larger rosette sizes (relative to the WT), *FTSHi3* overexpression lines had a much higher survival rate after 15 days of drought (Table S1). Most rosettes of the overexpression lines were free of drought symptoms (Figure 3, Table S1), similar to *ftshi3-1* (Mishra et al., 2021). While the amount of *FtsHi3* within the plant therefore seems to be directly correlated to plant growth, drought tolerance is afforded by deviations from WT amounts of *FtsHi3*. Similarly, *Zat10/STZ* has previously been identified as both a positive and a negative regulator of drought response in Arabidopsis (Mittler et al., 2006). However, whether FTSHi3 plays a role in positive or negative regulation of drought stress tolerance remains to be determined.

*pFtsHi3-OE* lines had significantly higher leaf water content, ranging from 1.1 to 1.6 times of WT following 15 days drought stress and three days rewatering (Figure 2D, - OE3 and -OE6 were somewhat lower). The positive water holding capacity of the overexpression lines under drought stress is likely a combination of several factors, including their reduced stomatal density and size (Figure S4A, B), leading to a higher WUE. Previously, targeted knockout of the *GTL1* (GT-2 LIKE 1) transcription factor, which represses STOMATAL DENSITY AND DISTRIBUTION1 (*SDD1*), was shown to reduce stomatal density without affecting photosynthetic efficiency (Yoo et al., 2010). The *gtl1* mutants as a result possessed greater WUE, similar to the *ftshi3-1* mutant line (Mishra et al., 2021). In addition, a recent comparison of over 300 Arabidopsis cultivars demonstrated that reduced stomatal size increases WUE in Arabidopsis in general (Dittberner et al., 2018). The combination of a reduction in these two traits clearly explains the increased WUE seen in our overexpression lines (Table S2).

Our ABA findings should also be explained through these physiological differences and the resulting increase in WUE. While ABA concentrations were similar in the WT and *pFtsHi3-OE1* and *-OE2* lines prior to drought, they were not elevated to the same extent post-drought (Figure S4C). Since accumulation of ABA induces guard cell closure (Daszkowska-Golec and Szarejko, 2013), and guard cell closure inherently improves WUE, reduced ABA levels (and thus open stomata) should result in decreased WUE. However, the opposite was observed in the *pFtsHi3-OE* and *35SFtsHi3-OE* lines (Table S2). This suggests that the *pFtsHi3-OE* lines are more sensitive to ABA, or that they are not correctly sensing drought. Interestingly, based on both our proteomics (Table S6) and qPCR data (Table S3), it appears that *ftshi3-1* and *pFtsHi3-OE1* are always perceiving drought to some extent, yet not so much as to manifest drought symptoms. This further suggests that FtsHi3 is at least partially involved in the communication of plant drought state to the chloroplasts.

### Pre-protein import related peptides accumulate in *ftshi3-1*

The *ftshi3* mutants possess a slow growing, chlorotic phenotype (Mishra et al., 2021), similar to that of the *hsp93/ clpC1* mutant lines (Sjögren et al., 2004). Mutations in *hsp93* cause dramatic pleiotropic effects that stymie plant development and chloroplast biogenesis. These mutant lines also show elevated amounts of other CLP protease sub-units (Sjögren et al., 2004). It is suggested that CLPC1 normally functions as a chaperone by acting as a translocation motor that binds directly to TIC110 (Akita et al., 1997). TIC110 is a core component in the TIC translocation complex and is responsible for the import of the majority of chloroplast targeted polypeptides (Kovacheva et al., 2005; Sjögren et al., 2014). Another chaperone protein that interacts with the TIC complex, HSP90-III/CR88, shows reduced stature and delayed chloroplast development associated with interrupted preprotein import (Cao et al., 2003). Interestingly, we observed an increase in the amount of HSP90, HSP93 and TIC110 in our proteomics data (as well as several other TICs and TOCs, Figure S6). Based on the compensatory mechanism observed in *hsp93* mutants (Sjögren et al., 2004) and the similar phenotypes observed in *ftshi3-1*, *hsp90* and *hsp93* (Cao et al., 2003; Sjögren et al., 2004; Mishra et al., 2021), it is possible that FtsHi3 plays a downstream role in pre-protein import into the thylakoid. This would be consistent with the growth-related phenotypes seen in the *ftshi3-1* mutant and *pFtsHi3-OE* lines, although generation of *ftshi3 hsp90* or *ftshi3 hsp93* double mutants should be carried out to verify this hypothesis.

The fully developed chloroplasts of *pFtsHi3-OE* lines were increased in width and more thylakoid membranes were observed than in the WT (Figure S2D-F). Similar findings were also observed in FtsHi1-YFP overexpression lines, which produced larger chloroplasts (Kadirjan-Kalbach et al., 2012). Since the corresponding *arc1* mutant produced smaller chloroplasts, that pseudo-protease was therefore suggested to negatively influence the timing of division.

FtsHi3 must be, at least in part, required for the transmission of growth promoting signals. This is further supported by the accumulation of proteins involved in isoprenoid metabolism, such as those for GGPP (Geranylgeranyl pyrophosphate) and FPP (farnesyl pyrophosphate). GGPP is a precursor to several phytohormones including gibberellins, strigolactones and abscisic acid (Lange and Ghassemian, 2003), whereas FPP is a substrate for the synthesis of a wide range of plant growth substances such as sterols, brassinosteroids and heme (Bouvier et al., 2005; Closa et al., 2010). The compensatory mechanisms for isoprenoids are unclear but given that these proteins are absent in the WT: *pFtsHi3-OE* comparison, one can conclude that interrupted growth signaling may result in an accumulation of growth promoting precursors.

### The *ftshi3-1* and *pFtsHi3-OE* lines have overlapping changes in peptide abundance

We sought to identify common responses in the *ftshi3-1* mutant and *pFtsHi3-OE* lines to explore their resistance to drought (Figure 5). A number of common proteins displayed opposing or differing abundances in *ftshi3-1* and *pFtsHi3-OE* lines (relative to the WT) under these conditions; KIN1, AK4RC9, the ABCG30, 36 and 37 as well as LEA4-5.

The ABCGs 30, 36 and 37 share a common identifying peptide, which makes an ABCG-specific discussion of their contribution to the drought resistance difficult. Nonetheless, that the ABCG protein group is elevated in both the *ftshi3-1* and *pFtsHi3-OE* lines, but repressed in the watered and droughted WT, respectively. ABCG30 has been demonstrated to actively import the stress hormone ABA into imbibed seeds, acting as a negative regulator of germination (Kang et al., 2015). Other work has demonstrated that *abcg30* mutants possessed different exudate profiles, which affected the bacterial and microbial communities around Arabidopsis roots (Badri et al., 2009). The combination of these suggests that ABCG30 may also regulate root ABA transport in Arabidopsis. ABCG36 is localized to the plasma membrane primarily in the root and leaf epidermis (Strader and Bartel, 2009). In addition to its well documented involvement as an indole-3-butyric acid transporter (Strader and Bartel, 2009), overexpression of *ABCG36* decreases the level of sodium *in planta* resulting in enhanced resistance to drought (Kim et al., 2010). This transporter also appears to utilize cadmium as a primary substrate, where it acts as an exclusion pump (Kim et al., 2007). Interestingly, *ABCG36* transcript accumulation does not appear to be induced by drought or salt stress, although it accumulates in response to cadmium via WRKY13 (Sheng et al., 2019). ABCG36 is also well documented to be involved in plant defense responses (Johansson et al., 2014), although, the relevance of this is not clear. *ABCG37* is plasma membrane localized and largely expressed in the root epidermis (Růžička et al., 2010). ABCG37 appears to share some substrate overlap with ABCG36 (Ziegler et al., 2017), but is reported to primarily function in the transport of Iron chelating coumarins to maintain iron nutrition (Fourcroy et al., 2014). The contribution of ABCG37 to drought is likely insignificant. It is possible that the elevated amounts of ABCG36 seen in both the *pFtsHi3-OE* and *ftshi3-1* lines may explain, at least in part, their drought resistance. The question of whether the corresponding transcripts are elevated remains unclear, although we were unable to detect a corresponding WRKY13 protein in our proteomics data. One possible explanation is that their elevated abundance is due to lower ABA levels seen in the OE lines, although the *ftshi3-1* mutant reportedly contains WT levels of ABA under drought (Mishra et al., 2021). Given the wide-ranging roles that these transporters play in plant development, as well as their involvement with many phytohormones, it is easy to comprehend why *ftshi3-1* mutants have such a striking phenotype. However, since ABA levels are reduced in the *pFtsHi3-OE,* and normal *ftshi3-1* lines, it is unlikely that ABCG30 is contributing to drought resistance.

LEA4-5 transcripts reach the highest levels in their family under stress conditions (Olvera-Carrillo et al. 2010). There are some post translational effects, as LEA4-5 protein abundance was higher in response to ABA than NaCl despite similar transcript levels. Interestingly, data from our *ftshi3-1* and *pFtsHi3-OE* lines appears contrary to that of Olvera-Carrillo et al. (2010), since in their analyses, reduced LEA4-5 transcription results in plants that have reduced biomass and fecundity and are drought sensitive. In kind, LEA4-5 overexpression lines are significantly drought resistant. In our dataset, peptide abundance of LEA4-5 is reduced in the *ftshi3-1* and *pFtsHi3-OE* lines, but these plants retain their drought resistance. Since the WT lines under drought accumulate the LEA4-5 peptide, as described by Olvera-Carrillo (2010), it is possible that the reduced abundance in the *ftshi3-1* and *pFtsHi3-OE* lines may be some sort of compensatory response.

The molecular function of KIN1 is unclear, but as a well-documented drought-and ABA-responsive element (Wang et al., 1995), our proteomics data shows a curious trend. In the WT under both drought and watered conditions, KIN1 is similarly abundant, whereas both the *ftshi3-1* and *pFtsHi3-OE* lines have markedly reduced abundance, in line with the idea that these lines are not perceiving drought. However, since the levels of ABA are highly elevated in the droughted WT, the drought responsiveness of KIN1 in our experiment is not so clear.

The ALDO-KETO REDUCTASE FAMILY 4 members (AKR4C8-C11) are responsible for the metabolism of various sugars, steroids and carbonyls. They are also implicated in the detoxification of harmful metabolites produced during stress. Of these AKR4C8 is suggested to possibly function in the metabolism of brassinosteroids (Simpson et al., 2009). Transcripts of AKR4C8 and AKR4C9 are both dramatically elevated under stress conditions (Simpson et al., 2009). In our dataset, AKR4C9 abundance is elevated in both *ftshi3-1* and *pFtsHi3-OE* lines. Under watered conditions, the WT has reduced abundance of AKR4C9 and in drought, the abundance of this enzyme is somewhat elevated, consistent with the drought responsiveness of AKR4C9 transcripts (Simpson et al., 2009). This effect is likely downstream of FtsHi3 and its exact role in conferring drought resistance remains unclear.

### Particular binding patterns recapitulate the non-monotonic behavior of putative FtsHi3 containing complexes

We used a mathematical approach to gain some insight into possible mechanisms that could give rise to our observations that the growth rate increases with increasing *FTSHi3* expression, yet stress tolerance shows non-monotonic behavior (Figure 6, Figure S8). A key aim of our modelling approach was to identify a minimal set of required features. We assumed the phenotype (either in terms of growth rate or stress tolerance) was determined by the proportion of specific pseudo-proteases within a population of interacting homo- or hetero-hexameric complexes. With this simple relationship, we found that complexes of at least three different interacting subunits could exhibit non-monotonic phenotypic behavior. The frequency of non-monotonic behavior increased in systems with four different interacting subunits, but it was not guaranteed. Indeed, our modelling results show that certain patterns of binding interactions are more likely to produce non-monotonic behavior. For example, in systems of 4 different (pseudo-) protease subunits, non-monotonic behavior is highest when FtsHi3 can bind to all other subunits. Moreover, in heteromeric complexes that show non-monotonic behavior, a (pseudo-) protease subunit does not bind to at least one of the other subunits. Thus, our modeling predicts that FtsHi3 should be able to bind multiple partners and some of its partners should not be able to bind to each other. We note that such predictions are the product of many simplifying assumptions made in the absence of precise mechanistic information and are likely to change, once more details are learned and included in the modeling. However, we conclude that non-monotonic phenotypic behavior should be expected in many systems of heteromeric complexes and does not require much mechanical complexity.

## Conclusion

The culmination of our findings strongly indicate that FtsHi3 plays a role in both growth and drought response. It is tempting to speculate that this pseudoprotease functions as an important switch that can inhibit growth when conditions are not favorable. Given the localization of this protein in the thylakoid membrane, it sits internal to the TIC/TOC preprotein import system. The idea that preproteins are further translocated via the putative FtsHi3 complex is not far-fetched as the other FtsHi proteins also possess this ability (Kikuchi et al., 2018; Schreier et al., 2018). It is further possible that a transient FtsHi3 complex is all that is required to act on the preprotein substrate. Very fast protein turnover would explain why FtsHi3 has yet to be detected in immunoprecipitation assays (Kikuchi et al., 2018) as well as why it was not detected in the *pFtsHi3-OE1* proteome.

In practice, we suggest that such a mechanism may exist that relies on the stoichiochemical balance of the FtsHi3 homo/ heterohexamer complex. In fact, substrate preference changes with altered complex composition has been demonstrated in other protease complex systems (Wojtkowiak et al., 1993). Under WT conditions, the hexamer exists in a state that allows both growth and drought signals to be communicated to the chloroplasts, finely controlling photosynthesis (i.e. growth vs drought resistance). In the *ftshi3-1* mutant, we see that neither growth nor drought signals are being perceived. However, in the overexpression lines, the complex exists in a state where substrate competition results in the transmission of putative growth signals, at the expense of as yet unknown drought signals. This system may also function in the inverse, but more biochemical studies are needed to explore this hypothesis.

## Material and methods

### Plant material and growth conditions

Arabidopsis wild type (WT) and mutant seeds were sterilized with 10% NaCl, washed 4X with sterile water, and then stratified for two days at 4°C. The seeds were selected on full-strength MS (Murashige & Skoog) agar (Murashige and Skoog, 1962), supplemented with 1% sucrose and 75 µg/l sulfadiazine. After 12 days post-germination on plates, the plants were transferred to soil. Water-deficit stress was applied on plants grown on soil in a growth chamber under short-day conditions (8 h/16 h photoperiod, 22°C/18°C), with a relative humidity of 50% and 150 μmol photons m^-2^ s^-1^ after approximately four weeks.

### Generation of transgenic Arabidopsis seedlings

To generate *FTSHi3* overexpression lines, genomic DNA fragments were amplified from *A. thaliana* by Phusion ® proofreading polymerase (Thermo Fisher Scientific USA). To generate plants with an additional *FTSHi3* gene under its native promoter (*pFtsHi3-OE* lines), a construct containing the amplified *FTSHi3* promoter sequence (as predicted by Knudsen, 1999) was generated using the primers ‘ftshi3 Promoter Forward’ and ‘ftshi3 Reverse for HA-line’ (Table S7). The *pAtftshi3::ftshi3* genomic DNA was cloned into a pENTR/D-TOPO vector and transferred into the destination vector pGWB15 resulting in a 3xHA tagged gene product. To generate plants overexpressing *FTSHi3* under a 35S promoter (*35SFtsHi3-OE*) the coding sequence of *FTSHi*3 was amplified and cloned into a pGWB5 vector under the control of the CaMV35S promoter, primers used are listed in (Table S7). The binary plasmids (*pAtFtsHi3::AtFtsHi3::HA* and *35S::AtFtsHi3::GFP*) were transformed into electro-competent *Agrobacterium tumefaciens*(GV3101::pMP90 (pTiC58DT-DNA); (Hellens et al., 2000). WT (Col-0) plants were transformed with these constructs by the floral dip method described by Clough and Bent (1998). For complementation (generation of *35SFtsHi3-comp* lines) *35S::AtFtsHi3::GFP* was transformed in the non-segregating T2 homozygous *ftshi3-1* T-DNA line of the GABI-KAT collection. The presence of the construct in the T1 and T2 generation was confirmed by germinating transgenic seeds on 35 mg/ml hygromycin-B selecting MS agar plates. The experiments were performed on T2 generation seeds.

### Water stress analysis

Wild-type and transgenic plants were grown on soil in the growth chamber under short-day conditions (8 h/16 h photoperiod, 22°C/18°C, relative humidity 50% and 120 μmol photons m^-2^ s^-1^) with regular watering for 3-4 weeks before treatment. Long-term water stress analyses were performed by withdrawing watering until the drought effects were observed in the respective genotypes. The experiments were performed three times.

### Drought tolerance assay

To assay drought tolerance, the ‘one rosette per-pot’ and ‘weighing’ methods were used as described by (Harb and Pereira, 2011; De Ollas et al., 2019). Two-week-old, plate-grown, WT and overexpression seedlings (15 replicates) were transplanted to 5 cm pots containing a 3:1 mixture of commercial soil (Hasselfors garden special, Sweden) and Vermiculite (Sibelco, Europe) and plants were grown in a growth chamber under short-day conditions (8 h/16 h photoperiod, 22°C/18°C). Initial daily transpiration was calculated on pots saturated with 500 ml water (15 pots per tray) and weighed for 5 to 6 days to determine the soil water volumetric content. Rosette diameters were measured at 3–4 day intervals throughout the experiment. Drought was induced at 37 days (23 days following transplantation) and pots were weighed daily between 9 and 11 AM to calculate water deficit. After 15 days of drought stress, plants were rewatered for three days, and rosette fresh weight (FW) was recorded. Rosettes were subsequently placed in glassine bags and oven-dried at 65 °C for 72 h to obtain dry weight, (DW). Leaf water content (%) was calculated as: (FW-DW/FW).

### Abscisic Acid Extraction and Quantification

Sample preparation and extraction for Solid Phase Extraction (SPE) and ultra-high performance liquid chromatography-mass spectrometry (UHPLC-MS/MS) were performed as described by Haas et al. (2021).

### Measurements of dehydration

The percentage of dehydration was determined by cutting and weighing well-watered plants as described by (Seo et al., 2000; Koiwai et al., 2004). Ten plants each of WT, *pFtsHi3-OE1 and pFtsHi3-OE2* at the age of four weeks were detached, and their fresh weight (FW) was measured. Left on the bench at RT, their weight was re-recorded after designated time intervals. The leaves’ relative water content was calculated by ((FW-Weight at any time point)/FW)*100).

### Phenotypic characterization

Wild type and mutant seedlings were investigated at the age of ten days using a Leica MZ9.5 stereomicroscope or scanned by Epson Perfection 3200 PHOTOscanner (Japan). Plants grown on soil or exposed to stress conditions were photographed at the age of six weeks using a Canon 650D camera.

### RNA extraction, cDNA synthesis, and quantitative PCR (qPCR)

RNA extraction and cDNA synthesis were performed according to (Mishra et al., 2019). The housing-keeping genes (ubiquitin, tubulin, and actin, Czechowski et al., 2005) and gene-specific qPCR primers are listed in Table S7.

### Leaf-level gas exchange

A portable photosynthesis system (Li-6400xt, Li-Cor, Lincoln, NE, USA) was used to determine the photosynthesis rate (assimilation *AN*) and stomatal conductance (*gs*) as described by (Tomeo and Rosenthal, 2018). Detailed steps are mentioned in (Mishra et al., 2021).

### Chloroplast size and ultrastructure

TEM was used to study the chloroplast morphology of the first true leaf of 12-day-old seedlings of WT. Sample preparation and microscopy were performed at the (UCEM) Umeå Centre Electron Microscopy. The open-source image-processing program ImageJ (Java-based image processing program developed at the NIH) measured chloroplast length and width.

### Mass spectrometry based proteomics

#### Sample preparation

Arabidopsis proteome samples were prepared from leaf tissue harvested from watered and drought treated WT, *ftshi3-1* and *pFtsHi3-OE* plants grown as mentioned in the drought tolerance section. Five replicates per genotype per treatment were pooled and five technical replicates were used. The harvested plant material (500 mg) was inserted in 2 ml Eppendorf tubes with glass beads and frozen in liquid N_2_. The tissue was homogenized using a TissueLyser at 30 Hz for 1 min and refrozen in N_2_. Proteins were extracted in 3 volumes of prechilled (−20°C) precipitation solution (TCA 10 % and Acetone 90 %) and precipitated at −20°C overnight. The proteins were pelleted by centrifugation at 14,800 g for 15 min at 4°C and supernatant was discarded. The pellet was then washed with 500 µl ice-cold acetone and centrifuged again as above. The washing step was repeated 3 x to remove residual TCA and acetone and the pellet was dried at room temperature and was stored at −80°C.

TCA-precipitated proteins were resuspended in 45 µl 8M Urea buffer (5 mM EDTA, 100 mM NH4HCO3, 1 mM DTT, pH 8.0). The total protein concentration was determined using the BCA protein assay kit (ThermoFisher Scientific). Subsequently, 15 µg total protein per sample was reduced (10 mM DTT, 30 min, 30 °C), carbamidomethylated (55 mM CAA, 30 min, room temperature) and digestion with trypsin (proteomics grade, Roche) overnight at 37°C at a 1:50 enzyme: protein ratio (w/w). Digests were acidified by addition of 0.5% (v/v) formic acid (FA) and desalted using self-packed StageTips (three disks per micro-column, ø 1.5mm, C18 material, 3M Empore). The peptide eluates were dried to completeness and stored at −80°C. For LC-MS/MS analysis all samples were re-suspended in 75 µl 2% acetonitrile (ACN) and 0.1% FA in HPLC grade water and 2 µl sample volume were injected into the mass spectrometer per mass spectrometric (MS) measurement.

LC-MS/MS data acquisition. LC-MS/MS measurements were performed on a nanoLC Ultra1D+ (Eksigent, Dublin, CA) coupled online to a Q-Exactive HF-X mass spectrometer (ThermoFisher Scientific). Peptides were loaded onto a trap column (ReproSil-pur C18-AQ, 5 μm, Dr. Maisch, 20 mm × 75 μm, self-packed) at a flow rate of 5 μl/min in 100% solvent A (0.1% FA in HPLC grade water). Subsequently, peptides were transferred to an analytical column (ReproSil Gold C18-AQ, 3 μm, Dr. Maisch, 400 mm × 75 μm, self-packed) and separated using a 50 min linear gradient from 4% to 32% of solvent B (0.1% FA in ACN and 5% (v/v) DMSO) at 300 nL/min flow rate. Both nanoLC solvents contained 5% (v/v) DMSO. The Q-Exactive HF-X was operated in data-dependent acquisition (DDA) mode, automatically switching between MS1 and MS2 spectrum acquisition. MS1 spectra were acquired over a mass-to-charge (m/z) range of 360–1300 m/z at a resolution of 60,000 using a maximum injection time of 45 ms and an AGC target value of 3e6. Up to 18 peptide precursors were isolated (isolation window 1.3 m/z, maximum injection time 25 ms, AGC value 1e5), fragmented by high-energy collision-induced dissociation (HCD) using 26% normalized collision energy and analyzed at a resolution of 15,000 with a scan range from 200 to 2000 m/z. Precursor ions that were singly charged, unassigned, or with charge states > 6+ were excluded. The dynamic exclusion duration of precursor ions was 25 s.

#### LC-MS/MS data analysis

Peptide and protein identification and quantification were performed using MaxQuant (version 1.6.17. 0) with its built-in search engine Andromeda (Cox et al., 2011; Tyanova et al., 2016). MS2 spectra were searched against an Arabidopsis (Arabidopsis Thaliana) proteome fasta file downloaded from TAIR (48359 protein entries, downloaded April 2017) supplemented with common contaminants (built-in option in MaxQuant). Carbamidomethylated cysteine was set as fixed modification and oxidation of methionine and N-terminal protein acetylation as variable modifications. Trypsin/P was specified as the proteolytic enzyme. The precursor tolerance was set to 4.5 ppm, and fragment ion tolerance to 20 ppm. Results were adjusted to 1 % false discovery rate (FDR) on peptide spectrum match (PSM) and protein level, employing a target-decoy approach using reversed protein sequences. The minimal peptide length was defined as 7 amino acids. The “match-between-run” function was disabled. Label-free protein quantification (LFQ vales) was used in downstream data analysis with the software Perseus (Tyanova et al., 2016). LFQ values were log-transformed (base 2) and median centred, followed by filtering proteins based on minimally one detected LFQ value throughout five biological replicates within in least one condition. Subsequently, imputation of missing values was carried out with Perseus using the function “replace missing values from normal distribution” (width = 0.3, downshift = 1.8, default settings). The functional annotation of proteins was downloaded from TAIR. The protein differential expression analyses were performed using the students t-test. The resulting p-values were corrected using the Benjamini and Hochberg method to control FDR (0.01) for differential expression. Gene ontological analysis were conducted using PANTHER (pantherdb.org) and string plots were generated using STRING (string-db.org). A list of significant AGIs from our comparisons can be found in the supplement (Table S8-10).

### Modelling methods

To receive clues in absence of mechanistic information about the different phenotypes arising with different amounts of FtsHi3 in the plant cell, we developed a mathematical approach that captured the essential features of protease heteromer-formation with a sampling-based exploration of model space.

We made assumptions based on observations of FtsH proteases in other membranes, in particular we assumed that FtsHi3 binds specific partners to form hetero-hexameric complexes (Moldavski et al., 2012). For simplicity, we assumed that each such heteromeric structure is formed by only two types of compounds in specific stoichiometric ratios of 1:1. In principle hexamers could be formed in many possible ways such as through sequential binding of proteases one at a time (Nauta and Miller, 2000) or through combinations of larger structures such as dimers or trimers (Prokhorova and Blow, 2000). If we considered all such modes of forming hexamers then the parameter space would increase significantly and our results would conflate the effects of binding interactions with the mechanism of hexamer formation. Thus, we made the simplifying assumption that hetero-hexamers are composed of two different subunits, which means that our model is applicable to any system, in which basic components are combined in pairs, e.g. monomers forming dimers or trimers forming hexamers. Without knowledge of which FtsH subunits bind to each other, we constructed a set of all possible interaction diagrams (17 different diagrams) for systems with 3 or 4 proteases that could result in heteromeric complexes. Determining the typical qualitative dynamics of each interaction diagram would identify protease binding interactions that could produce the experimental observations. We determined the typical qualitative dynamics of an interaction diagram by studying its corresponding system of differential equations. Each interaction diagram of 3 or 4 proteases can be translated into a system of 3 or 6 differential equations, respectively. In the absence of information concerning the reaction kinetics of binding, we determined the typical behavior through a sampling procedure. We randomly sampled reaction rates using a uniform distribution between (0,1) and initial protease concentrations using a uniform distribution between (0,10). For each random set of parameters, we solved the differential equation system to find the steady state concentrations of hexameric complexes. We then used subunit A as a proxy for FtsHi3 and repeated the steady state calculation after increasing/decreasing its initial concentration by a factor of 100, representing the effects of overexpression/knock downs. We assessed whether increasing the concentration of FtsHi3 (or A) resulted in: 1. an increased proportion of steady state concentration of A complexes, and 2. non-monotonic behavior in the proportion of complexes containing any other protease. The requirement for non-monotonic behavior was either that the proportion of complexes formed by a protease should be high-low-high or low-high-low with increasing concentration of subunit A. We calculated the average occurrence of each qualitative behavior for an interaction diagram using 10 samples of 100 sets of random parameters.

### Data availability

All mass spectrometric raw files as well as the MaxQuant output files have been deposited to the ProteomeXchange Consortium (http://proteomecentral.proteomexchange.org) via the PRIDE partner repository with the dataset identifier PXD037172 (reviewer account details: Username; Password 3utDe55B).

## Acknowledgements

This work was supported by grants from the Knut and Alice Wallenberg Foundation (KAW 2016.0341 and KAW 2016.0352) and the Swedish Governmental Agency for Innovation Systems (VINNOVA, 2016-00504). We acknowledge financial support by the Swedish Research Council VR (grant number 2019-04472) and Umeå University. We are grateful for the help of the KBC electron microscopy platform and the Swedish Metabolomics Centre supported by Umeå University and the Swedish University of Agricultural Sciences. This work has been supported by EPIC-XS, project number 823839, funded by the Horizon 2020 programme of the European Union. We would also like to thank the Kempe foundation for the postdoctoral fellowship awarded to SDC.

## Author Contributions

Conceptualization: LSM, CF; Investigation: LSM, SDC, SM, HI; Methodology: LSM, SDC, HI, EL; Data Curation: LSM, SDC, IS, MA, CL, HI, EL; Writing original draft: LSM, SDC, CF; Review and editing: all authors. Visualization: LSM, SDC, HI; Project administration and funding acquisition: CF.

**S. Figure 1. A,** Phenotypes of WT and *35SFtsHi3-OE5* to *-OE8* at 6 weeks of age under SD conditions (8/16 hr D/N cycle, 22°C). **B,** Rosette diameter of WT and *35SFtsHi3-OE5* to *-OE8* grown under the same conditions as **A** (n=8). **C,** Expression of *FtsHi3* relative to UBQ5 and Actin for each of the *35SFtsHi3* lines at 3 weeks of growth under the same conditions as **A** (n=3). Data are means ± standard error, asterisks indicate significance; p<0.05 (*), p< 0.01 (**), p< 0.001 (***) and scale bars are as indicated.

**S. Figure 2. A**, Root phenotype of 8-day-old seedlings from WT, *pFtsHi3-OE1* and *pFtsHi3-OE2* under SD conditions (8/16 hr D/N cycle, 22°C). Arrows indicate the lateral roots. **B,** Root length in 8-day-old seedlings (n=25). **C,** Number of lateral roots in 8-day-old seedlings (n=21). **D**, Transmission electron micrographs showing chloroplasts of the first true leaves in seedlings of WT, *pFtsHi3-OE1* and *pFtsHi3-OE2* (upper), and zoomed in ultrastructure showing grana stacking (lower), scale bars are 1μm. **E,** Size of chloroplasts from first true leaves in WT, *pFtsHi3-OE1* and *pFtsHi3-OE2* lines grown under the same conditions as **A** (n=40). **F,** Number of grana stacks per chloroplast under the same conditions as **A** (n=40). Data are means ± standard error, significances as per **S. Figure 1**, scale bars are as described.

**S. Figure 3**. Effect of drought on WT and *35SFtsHi3-OE5* to *OE8* lines grown in a growth chamber under SD, soil grown conditions (8/16 hr D/N cycle, 22°C). **A,** Rosette diameter prior to, and following, drought induction at day 37. **B,** Progressive % water deficit during drought. **C,** Dry weight of harvested rosettes following drought. **D,** Calculated leaf water content %: (Biomass of watered control) - (Biomass of droughted) / (Biomass of watered control). Data are means ± standard error (n=15), significances as per Figure 1.

**S. Figure 4.** Stomatal physiology and ABA quantification in WT, *pFtsHi3-OE1* and *- OE2* lines. **A,** Reduced stomatal size (μm) and **B,** Stomatal density (number per mm^2^) in *pFtsHi3-OE* lines (n=5, 30 stomata per plant). **C,** Quantification of endogenous ABA levels (ng g^-1^ FW) in watered and drought-stressed leaves of WT, *pFtsHi3-OE1* and *pFtsHi3-OE2* (n=4). **D,** Representative microscopic images of stomatal response to treatment with 10 µM ABA. **E,** Stomatal aperture size in response to treatment with 10 µM ABA (n=5, 45 stomata per plant). Data are means ± standard error, significances as per Figure 1. Scale bars, as described.

**S. Figure 5.** Volcano plots showing number of proteins with statistically altered abundance in **A**, *ftshi3-1* vs WT under drought conditions. **B,** *ftshi3-1* vs WT under watered conditions. **C,** *35SFtsHi3-OE* vs WT under drought conditions. **D,** *35SFtsHi3-OE* vs WT under watered conditions. Data are log_2_ Fold Change (x-axis) vs −log_10_ of the corresponding *p*-value, with an S0 cutoff of 2 and an FDR cutoff of 0.01 (regression lines). **E,** Venn Diagram showing the number of overlapping differentially abundant proteins in *ftshi3-1* (vs WT) under drought (A) and watered (B) conditions and their relationship to (C) *35SFtsHi3-OE* (vs WT). Overlapping fields BC are indicated by * and ABC by †.

**S. Figure 6.** STRING plot showing protein: protein interactions between **elevated** proteins in *ftshi3-1* relative to WT under drought conditions. Data were thinned to retain only high confidence interactions (0.700) and disconnected nodes were removed. The resulting elements were clustered using an MCL inflation parameter of 2. Clusters are distinguished by node color and line color indicates number and type of evidence (see string-db.org/), dashed lines indicate borders of clusters. Peptide names or AGIs are provided adjacent to nodes.

**S. Figure 7.** Histogram showing average log_10_ transformed LFQ values and their corresponding standard deviation for the greater FtsH protein family. Colors represent different contrasts, *ftshi3-1* drought (Dark blue), *ftshi3-1* watered (Orange), *pFtsHi3-OE* drought (Grey), *pFtsHi3-OE* watered (Yellow), WT drought (Light Blue), WT watered (Green). Subcellular localization of the AtFtsH members is given along the X-axis. ‘a’ indicates measurements based off a single measurement.

**S. Figure 8.** Behavior profile of FtsHi3 interactions with other FtsH/FtsHi subunits in a hexamer. Bar graphs with the proportion of samples (the mean of 10 samples of 100 simulations) that have monotonic behavior in interactions between subunit A (e.g. FtsHi3; red) and non-montonic behavior in other subunits (e.g. (pseudo-) proteases; blue). All interactions allow monotonic behavior for subunit A except those in panels B) and L), where the proportion of A complexes was 1 for each line (WT, *ftshi3-1* and *pFtsHi3-OE*). Non-monotonic behavior was more frequently detected in interactions where subunit A can interact with more than one other compound.

## Tables

**S. Table 1.** Evaluation of *pFtsHi3-OE and 35SFtsHi3-OE* lines compared to WT following 15 days of drought

**S. Table 2.** Gas-exchange properties of WT*, pFtsHi3-OE1*-*2* and *35SFtsHi3-OE5,7,8* lines grown in watered or drought conditions. Values are net photosynthesis (*A_N_*, µmol CO_2_ m^-2^s^-1^) and stomatal conductance (*gs*, mol CO_2_ m^-2^s^-1^), CO_2_ concentration inside the leaf (*Ci*, µmol CO^2^ mol^-1^ air) and intrinsic water use efficiency (*WUE*). Data are means ± standard error, significances as per **S. Figure 1** caption.

**S. Table 3.** Relative expression of dehydration-induced and ABA-responsive genes in WT, *pFtsHi3-OE1* and *pFtsHi3-OE2* plants under watered or drought conditions. Data are normalized to the expression of *actin*. Data are means ± standard error, significances as per **S. Figure 1** caption

**S. Table 4.** Major gene ontological processes in elevated AGIs from *ftshi3-1* lines (vs WT) under drought conditions

**S. Table 5.** Major gene ontological processes in repressed AGIs from *ftshi3-1* lines (vs WT) under drought conditions

**S. Table 6.** Major gene ontological processes altered in both *pFtsHi3-OE* and *ftshi3-1* lines (vs WT) under drought conditions

**S. Table 7.** Names and sequences of primers used in this study

**S. Table 8.** List of significant AGIs in the ftshi3-1 vs WT contrast under watered conditions

**S. Table 9.** List of significant AGIs in the ftshi3-1 vs WT contrast under drought conditions

**S. Table 10.** List of significant AGIs in the pFtsHi3-1 vs WT contrast under drought condition

